# *Streptococcus pyogenes* infection and the human proteome with a special focus on the IgG-cleaving enzyme IdeS

**DOI:** 10.1101/214890

**Authors:** Christofer A. Q. Karlsson, Sofia Järnum, Lena Winstedt, Christian Kjellman, Lars Björck, Adam Linder, Johan A. Malmström

**Affiliations:** Lund University, Division of Infection Medicine, Department of Clinical Sciences, Lund, Sweden; Hansa Medical AB, Lund, Sweden

**Keywords:** Human proteome, Infectious diseases, Systems biology, Host-bacteria interactions, Mass spectrometry

## Abstract

Infectious diseases are characterized by a complex interplay between host and pathogen, but how these interactions impact the host proteome is unclear. Here we applied a novel mass spectrometry based proteomics strategy to investigate how the human proteome is transiently modified by the pathogen *Streptococcus pyogenes*, with a particular focus on bacterial cleavage of IgG *in vivo*. In invasive diseases, *S. pyogenes* evokes a massive host response in blood, whereas superficial diseases are characterized by a local leakage of several blood plasma proteins at the site of infection including IgG. *S. pyogenes* produces IdeS, a protease cleaving IgG in the lower hinge region and we find highly effective IdeS-cleavage of IgG in samples from local IgG poor microenvironments. The results show that IdeS contributes to the adaptation of *S. pyogenes* to its normal ecological niches. Additionally, the work identifies novel clinical opportunities for *in vivo* pathogen detection.

## Introduction

The molecular interplay between host and pathogen is complex and under considerable evolutionary pressure. Determining the structure, function and dynamics of all host-pathogen interactions in a systems-wide and quantitative manner is a formidable and fundamental challenge in medical microbiology. A complicating factor is that many bacterial pathogens can cause diverse clinical manifestations, ranging from superficial to invasive disease, expanding the types of interactions. Commonly, local manifestations are confined to highly specific microenvironments like for example mucous layers and epithelial surfaces. In contrast, the development of invasive diseases can result in system-wide spread via the blood stream exposing the disease-causing pathogen to microenvironments associated with high concentrations of antibodies, complement proteins and professional phagocytes. Recent large-scale inventories of the human proteome have revealed that the molecular compositions of common infection sites are highly distinct under healthy conditions (Kim et al., 2014; Uhlén et al., 2010; Wilhelm et al., 2014). However, during an infection the molecular composition of these infection sites can be distorted by for example vascular leakage, activation of innate and adaptive immunity and through the infiltration of inflammatory cells. For example, previous large-scale quantitative analysis of the blood plasma proteome in an animal model of sepsis revealed that the blood plasma proteome is profoundly reorganized depending on disease severity (E. Malmström et al., 2016). In most cases, however, it is unknown whether molecular mechanisms altering the host proteome during an infection are a result of an active host defense or if some of the proteome changes are caused specifically by the infecting pathogen.

In this work, we hypothesized that the transient modulation of the human proteome by bacterial pathogens depends on the state of the host proteome at the infection site and the nature of the virulence factor repertoire of the infecting pathogen. To test this hypothesis, we selected the important bacterial human pathogen *S. pyogenes* as our model system since several molecular mechanisms of virulence are well characterized with an extensive list of confirmed and suggested virulence factors (Bisno et al., 2003; Cole et al., 2011; Cunningham, 2000; Musser and Shelburne, 2009). *S. pyogenes* infections cause diverse clinical manifestations ranging from mild and common local infections such as tonsillitis, impetigo and erysipelas to life-threating systemic disease like sepsis, meningitis or necrotizing fasciitis. Notably, the incidence of these diseases differs greatly, with more than 600 million cases of tonsillitis and less than 200,000 cases of lethal invasive disease annually (Carapetis et al., 2005). As mentioned, these large differences indicate that microenvironments associated with high disease incidence may represent natural habitats where the virulence factor repertoires have evolved to execute their primary functions (Wollein Waldetoft and Råberg, 2014). Consequently, diminishing effective protection by *S. pyogenes* virulence factors in other microenvironments is a possible explanation why cases of severe invasive infections are rare compared to common superficial and uncomplicated infections.

*S. pyogenes* has several independent virulence mechanisms that modulate humoral immunity. For example, the cell wall anchored surface proteins M (Fischetti, 1989) and H have domains that support protein-protein interactions with human host proteins to inhibit phagocytosis, promote adhesion and to secure nutrients from the host (Carlsson et al., 2003; J. A. Malmström et al., 2011). In addition, these proteins mediate nonimmune Fc-mediated binding of IgG antibodies resulting in evasion of phagocytosis (Åkesson et al., 1994; Heath and Cleary, 1989), mostly relevant in IgG-poor environments (Nordenfelt et al., 2012). Furthermore, *S. pyogenes* secrets enzymes associated with different mechanisms to prevent the normal function of IgG. EndoS hydrolyzes the conserved N-linked glycan on the heavy chains of human IgG (Collin and Olsén, 2001), resulting in a reduced affinity to most IgG-Fc receptors (Allhorn et al., 2008). SpeB is an unspecific protease capable of cleaving several proteins among them IgG (Collin et al., 2002). In addition, *S. pyogenes* secretes IdeS, a cysteine protease that specifically cleaves all four subclasses of human IgG in the lower hinge region of the heavy chain, generating one F(ab’)2 fragment retaining full antigen-binding activity and one non-covalently linked dimeric Fc fragment (Pawel-Rammingen et al., 2002a). As a consequence, IgG antibodies that are bound to the bacterial surface and cleaved by IdeS will lack IgG-Fc receptor and complement binding/activation capability. Apart from implications as an important anti-phagocytic virulence factor (Pawel-Rammingen et al., 2002b), IdeS is being developed as a biopharmaceutical drug to temporarily remove pathogenic IgG in autoimmune conditions (Collin and Björck, 2017), and is also currently undergoing clinical trials in kidney transplant patients to evaluate its capacity to remove pathogenic donor-specific antibodies and make highly sensitized patients eligible for transplantation (S. C. Jordan et al., 2017; Winstedt et al., 2015).

So far, the mode of action for many *S. pyogenes* virulence factors has been determined *in vitro* or in animal studies, whereas few mechanisms have been confirmed to take place in the human host. It can be anticipated that the combined *in vivo* activity of the *S. pyogenes* virulence factor repertoire will introduce differential abundance changes in the human proteome in addition to protein cleavage of both bacterial and human proteins. In this work we have developed novel targeted quantitative mass spectrometry methods to determine the impact of *S. pyogenes* infections on the human proteome with a particular focus on IgG cleavage at different infection sites. We show that invasive disease caused by *S. pyogenes* results in a systemic molecular host response in the blood plasma, which resembles host responses against other bacterial pathogens. In contrast, *S. pyogenes* causes a highly specific modification of the human proteome through IdeS-mediated proteolytic processing of IgG, which is most pronounced in non-invasive diseases. The results demonstrate that the combined *in vivo* effect of IgG cleavage by IdeS is dependent on the microenvironments associated with different disease and infection types. We anticipate that the proteomics strategy developed and described here can be extended to diseases caused by other bacterial pathogens to investigate the effect of individual virulence factors on host proteomes in patient samples.

## Results

### Development of an SRM-MS assay to measure the proteolytic activity of IdeS

Measuring the proteolytic effect by *S. pyogenes* on the entire IgG pool in complex clinical samples, requires new experimental assays to accurately monitor the relationship between intact and fragmented IgG at the subclass level. In this context, it is important to quantify several factors that may influence the degree of IgG cleavage, i.e. absolute amounts (mg/ml) of the IgG subclasses, IdeS, plasma proteins of relevance, and quantitative information of other proteases capable of cleaving IgG such as SpeB (Figure 1A). Since the assessment of IdeS activity entails analysis of samples where the absence is equally important as the presence of a signal, selected reaction monitoring mass spectrometry (SRM-MS) represents a suitable technique. For the construction of an absolute quantification method for IdeS using SRM-MS, we selected crude IdeS SRM-MS peptide-assays (Karlsson et al., 2012) and tested the performance of the crude assays in a dilution series of recombinant IdeS spiked in human serum (Figure S1). Based on these results, we selected two refined SRM-MS peptide assays (Figure 1B). For these two peptides we synthesized heavy labeled reference peptides to allow absolute protein quantification. As outlined in Figure 1A, we also included highly refined SRM-MS peptide assays for selected plasma proteins (Sjöholm et al., 2014) and a previously developed SRM-MS method to assess the number of *S. pyogenes* cells (Sjöholm et al., 2017). In this way it becomes possible to correlate the number of *S. pyogenes* cells to the IgG concentration and the degree of IgG breakdown.

**Figure 1.**
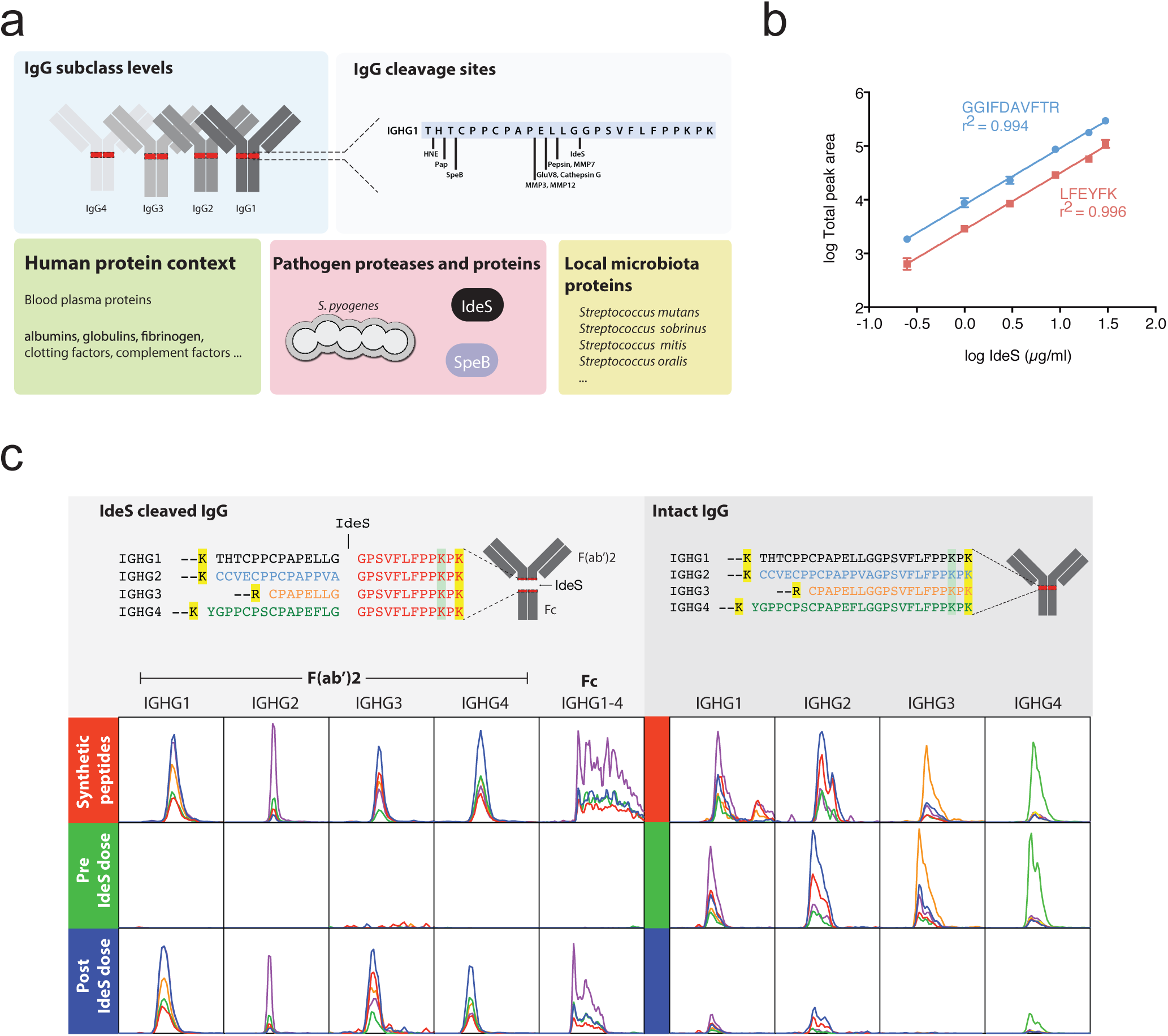
Development of SRM assays targeting the proteolytic activity of IdeS on IgG. A) Overview of targeted protein types used for the development of a quantitative SRM assays method to quantify the degree of proteolytic processing of IgG. The SRM method includes all potential IgG peptides produced by the targeted proteases in combination with trypsin. Measuring species-specific pathogen peptides requires knowledge of peptide specificity and sensitivity across relevant taxa. B) Six samples of IdeS at different concentrations in two different human serum backgrounds were prepared by serial dilution to generate a calibration curve spanning a concentration range of 0.25 μg to 30 μg IdeS per ml human serum. The plot shows the total area over the dilution range of indicated SRM targeted peptides derived from IdeS and their respective correlation coefficients (r^2^). The data represents average total area ± SD of the two different serum backgrounds. C) IdeS cleavage of IgG results in the formation of Fc and F(ab)2 fragments. Tryptic peptides derived from intact IgG or IdeS-cleaved IgG in the lower hinge region spanning the IdeS specific cleavage site in all four subclass heavy chains (UniProt accession entry / residue positions): IGHG1_HUMAN/104-137, IGHG2_HUMAN/101-134, IGHG3_HUMAN/158-185 and IGHG4_HUMAN/101-135) are shown in the upper panels. Yellow or green marked residues indicate a C-terminal trypsin cleavage site or a low probability C-terminal trypsin cleavage site, respectively. The peptide text color indicates the subclass specificity: black IGHG1, blue IGHG2, orange IGHG3, green IGHG4, redIGHG1-4. Based on synthetic variants of these peptides, SRM assays were developed. In the lower panels are comparisons of the SRM traces from the synthetic peptides (red) and human serum pre- (green) and post- (blue) intravenous administration of IdeS. The x-axis shows the normalized peptide retention times (iRT score). The y-axis’s ranges are fixed between pre- and post-IdeS doses to show level of change. The line colors represent different product ions depending on the peptide precursor.

For the construction of SRM-MS peptide assays for all four IgG subclass heavy chains we relied on conserved amino acid sequences as well as the major sequence variations found in the hinge region (Figure 1C). We used both the conserved and variable sequence identities to develop specific SRM-MS peptide assays to quantify total IgG and the individual IgG subclasses (Hong et al., 2013; Nordenfelt et al., 2012). These SRM-MS IgG peptide assays with associated heavy labeled reference AQUA peptides targets peptides in the Fc or F(ab’)2 regions of the IgG heavy chain, and allows the absolute quantification of the four IgG subclasses as previously shown (Nordenfelt et al., 2012). IdeS is a uniquely specific enzyme cleaving all subclasses of IgG in the lower hinge region (Pawel-Rammingen et al., 2002a). To develop SRM-MS peptide assays for determining the ratio between IdeS cleaved and intact IgG, we aligned the lower hinge region sequences spanning the IdeS P1-P1’ cleavage sites (Pawel-Rammingen et al., 2002a; Wenig et al., 2004) of the four subclasses heavy chains, and selected the peptides generated by trypsin cleavage alone and after a combination of both IdeS and trypsin (Figure 1C). Tryptic peptides spanning the IdeS cleavage site were unique for all four subclasses and represent suitable proxies for measuring the amount of intact IgG. Sequential cleavage of IgG using IdeS and trypsin generates unique N-terminal and C-terminal peptide fragments that are identical for all sub-classes (Figure 1C). We targeted all these peptides for the development of SRM-MS assays together with a set of negative control peptides with single negative or positive offset residuals (P2-P1 and P1’-P2’), which are hypothetically not generated by the IdeS and trypsin cleavage combination. As further controls, we also included peptides generated by other known specific proteolytic processing of IgG1 (Brezski and R. E. Jordan, 2010) in combination with trypsin digestion as seen in Figure 1A. For more information regarding the development of the SRM-MS assay see Table S2 and Figures S2-S4.

### Quantification of the proportions between IdeS cleaved IgG and intact IgG

Next we used the developed SRM-MS assay to test its specificity and quantitative performance to measure IgG degradation by IdeS both *in vitro* and *in vivo*. For *in vitro* testing, we used purified human monoclonal IgG antibodies (mAb) of subclass 1-4 and benchmarked the SRM-MS assays by side-by-side comparison with non-reducing SDS-PAGE (Figure S2). The results reveal a sequential cleavage of all subclasses with a rapid generation of single cleaved IgG (scIgG) and a slower generation of fully digested IgG (F(ab’)2 fragment) as previously reported (Vindebro et al., 2013). IgG3 is cleaved faster compared to the other subclasses and was fully digested after only 10 min of IdeS incubation (Figure S2).

To determine the capability of IdeS to cleave IgG in serum provided that sufficient amount of IdeS is available, we used 67 serum samples from a clinical phase I trial (Winstedt et al., 2015) (Table S1). In this trial twenty subjects were infused with 0.01, 0.04, 0.12 or 0.24 mg IdeS per kg bodyweight over 14 or 29 minutes. The lowest dose corresponds to a theoretical IdeS concentration of ~7 nM (~250 ng/ml, 89.25 aM on column), and is five orders of magnitude lower compared to the levels of human serum albumin (634 μM). This difference in dynamic range is just within the theoretical range of SRM-MS using un-fractioned serum or plasma (Mitchell, 2010). To determine IdeS pharmacokinetics, serum samples of the subjects were collected pre-treatment, directly after administration and at eight time points up to seven days. First we analyzed the serum concentration of IdeS in these samples starting from the first time points after administration using the optimized SRM-MS peptide assays. Overall the relationship between administered IdeS dose and the serum IdeS concentration was linear although showing an individual variation (Figure 2A). The measured IdeS concentrations depending on dose were in general in agreement with the theoretical values of IdeS serum levels (see Figure 2A) estimated from 3.8% serum per kg bodyweight (BW) (Morgan, 2005).

**Figure 2.**
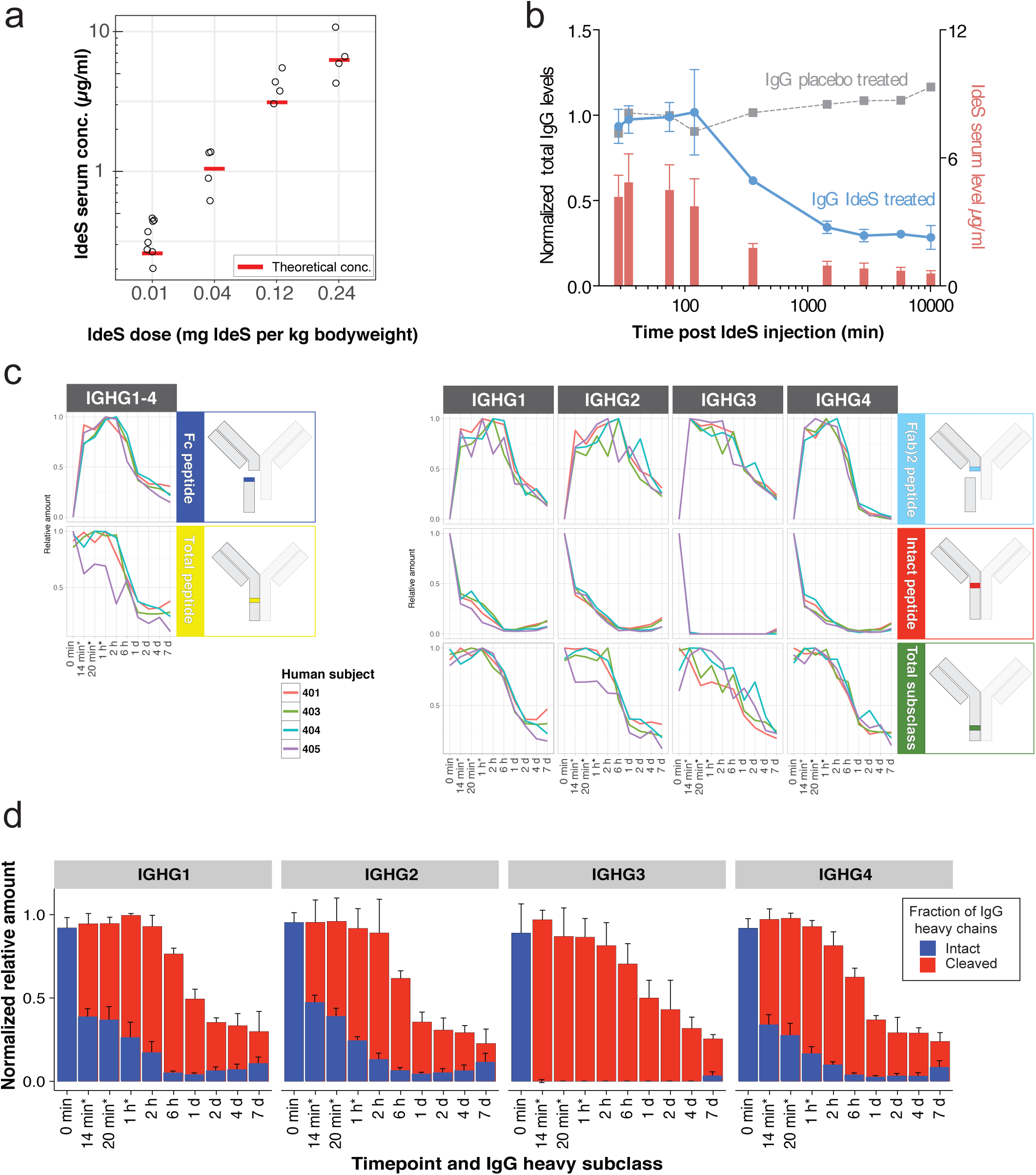
Time course of IdeS-cleavage of human IgG *in vivo* measured with SRM. A) The peak serum concentration (left y-axis) of IdeS across different administered IdeS doses in humans determined with SRM (black circles) and the theoretical serum concentration (red boxes). B) The plot shows the average total IgG heavy chain levels (sum of intact and IdeS-cleaved IgG based on peptide DTMSLR intensities normalized to total IgG before IdeS administration, blue, left y-axis) and IdeS (red, right y-axis) levels ± SD in four individuals treated with 0.12 mg IdeS per kg bodyweight over the time course period. The total IgG of a control individual injected with buffer is shown in grey. C) Peptide intensities of total IgG and subclass specific peptide levels of intact and IdeS generated tryptic peptides in four different human subjects over time. Schematic inserts to the right indicate location of the measured peptides. Asterisk marked time points (*) are as follows for subject 401 only: 29 min, 35 min and 75 min respectively. D) Average estimated ± SD levels of intact and IdeS-cleaved IgG at the subclass level after IdeS administration are calculated from measurements in C) (subjects 401, 403-405). Data are normalized to IgG subclass levels before IdeS administration. Asterisk marked time points (*) indicate slightly different time points for subject 401

In the same experiment, we quantified the amount of IgG in the time-course samples. Over the time-course experiment there was a clear reduction of the total IgG levels (here defined as the sum of intact IgG and IgG fragments) down to approximately 40% of the original levels in subjects treated with 0.12 12 mg/kg BW (Figure 2B and Table S3). In contrast, and as expected, the levels of IgG in the placebo treated control samples were not affected (Figure 2B). These results indicate that one major effect of intravenous IdeS administration to humans is a reduction of the total IgG levels. This was also confirmed in the phase I trial study using other methods (Winstedt et al., 2015). In the same experiments we also measured the level of intact and IdeS cleaved IgG reporter peptides. As shown in Figure 2C, the IdeS truncated subclass F(ab’)_2_-peptides, representing cleaved IgG, rapidly peaks within the first hour of IdeS administration after which the signal gradually declines. An example of the obtained results with the SRM-MS assays is shown in Figure 1C where targeted peptides show a reverse relationship to the signals from the peptides targeting intact and IdeS cleaved IgG (Figure 1A). Conversely, the intact IgG peptides spanning the cleavage site reduce rapidly in intensity for all the four subclasses with a more pronounced reduction of the IgG3 specific intact peptides. Interestingly, there are only minor differences between the healthy subjects included in the study (Figures 2C and S3). By correlating the signal from the subclass specific F(ab’)_2_-peptides to the intact IgG peptides spanning the cleavage site, it is possible to calculate the degree of cleavage across all the investigated patients (Figures 2D and S3). These results show that IdeS consistently increases the ratio between cleaved and intact IgG up to six hours after IdeS administration, to a point where >95% of the IgG heavy chains are cleaved, resulting in a completely disrupted IgG pool consisting of Fc, F(ab’)_2_ and scIgG fragments. The 6-hour time point correlates with a starting decline in the measured levels of IdeS in these samples, and with the overall reduction of the concentration of IgG in these samples. Six hours after IdeS administration, there is only a gradual increase of intact IgG over the tested seven days period. Again, IgG1, 2 and 4 display a similar degree of degradation whereas IgG3 is more sensible to IdeS cleavage. To estimate the accuracy of the signal from the peptides targeting IdeS activity we searched for peptides potentially generated by offset IdeS cleavage site (non P1-P1’ positions), peptides generated by other proteases than IdeS where the known cleavage sites for IgG1 were obtained from (Brezski and R. E. Jordan, 2010), and low probability tryptic peptides variants of the peptides targeting IdeS activity; in this latter case peptides with C-termini lacking the PK residues (Figure S4). In these experiments we note that these unexpected peptides account for less than 10% of the total signal, and hence that the estimated ratio between cleaved and noncleaved IdeS peptides is within this accuracy range. Collectively, these results demonstrate that IdeS reduces the entire pool of intact IgG up to at least 95% in healthy individuals provided that sufficient amount of IdeS is present.

### Proteomics analysis of host response during *S. pyogenes* infections

*S. pyogenes* causes a wide range of diseases associated with distinct host microenvironments. However, it currently remains unclear how these microenvironments are altered during an infection and how the specific host-proteomes influence cleavage efficiency of IgG by bacterial proteases. To determine the impact of *S. pyogenes* infections on the human proteome, we swabbed patients with tonsillitis, representing mild and superficial disease, and healthy controls. We also sampled blood plasma from sepsis patients with positive cultures for *S. pyogenes* or other bacterial pathogens, and wound fluid from patients with necrotizing fasciitis, representing invasive and severe disease (Figure 3A and Table S1). All samples were digested with trypsin and analyzed using data-independent acquisition mass spectrometry analysis (DIA-MS). The unsupervised hierarchical clustering of the DIA-MS data revealed two distinct sample clusters (Figure 3B). The first sample cluster is comprised of all plasma samples plus three of the wound fluid samples, whereas the second cluster contains all tonsil swabs and the remaining wound fluid samples. As shown in Figure 3B, the two sample clusters have distinctly different proteomes subdivided by two predominant protein clusters. Gene ontology (GO) overrepresentation test identifies an enrichment of intracellular proteins such as cytoskeletal, organelle, nucleus and cytosol proteins in protein cluster 1, and extracellular proteins, including blood plasma and vesicle lumen proteins in protein cluster 2. Pairwise comparison between plasma samples from the healthy controls and all sepsis patients, independent of bacterial pathogen, reveal 171 significantly induced or repressed proteins after correction for multiple hypothesis testing (Figures 3C and 3E). The most pronounced change was the increased levels of several acute phase proteins, such as serum amyloid A-1, serum amyloid A-2 and c-reactive protein (CRP)(Davidson, 2013), all over 500-fold more abundant in the sepsis patients (Table S4). On the other extreme, we observe a significant 20-100 fold reduction of several tissue derived proteins such as Myosin-9, ATP synthase subunit beta and 14-3-3 protein gamma, in addition to several known negative acute phase proteins like serum albumin and serotransferrin (Davidson, 2013; Ritchie et al., 1999) (31 and 35% reduction respectively). These results show that sepsis drastically distorts the composition of the blood plasma proteome as previously shown in animal models (E. Malmström et al., 2016). To test if the observed differences were specific for *S. pyogenes*, we performed a pairwise comparison between the sepsis plasma samples positive for *S. pyogenes* and the samples from sepsis patients infected with other bacterial pathogens. This comparison revealed relatively few statistically induced or repressed proteins, although the fold-change for some proteins was substantial (Figure 3F). The large non-significant protein fold abundance changes indicate a considerable heterogeneity in host response to sepsis, which is unrelated to pathogen type.

**Figure 3.**
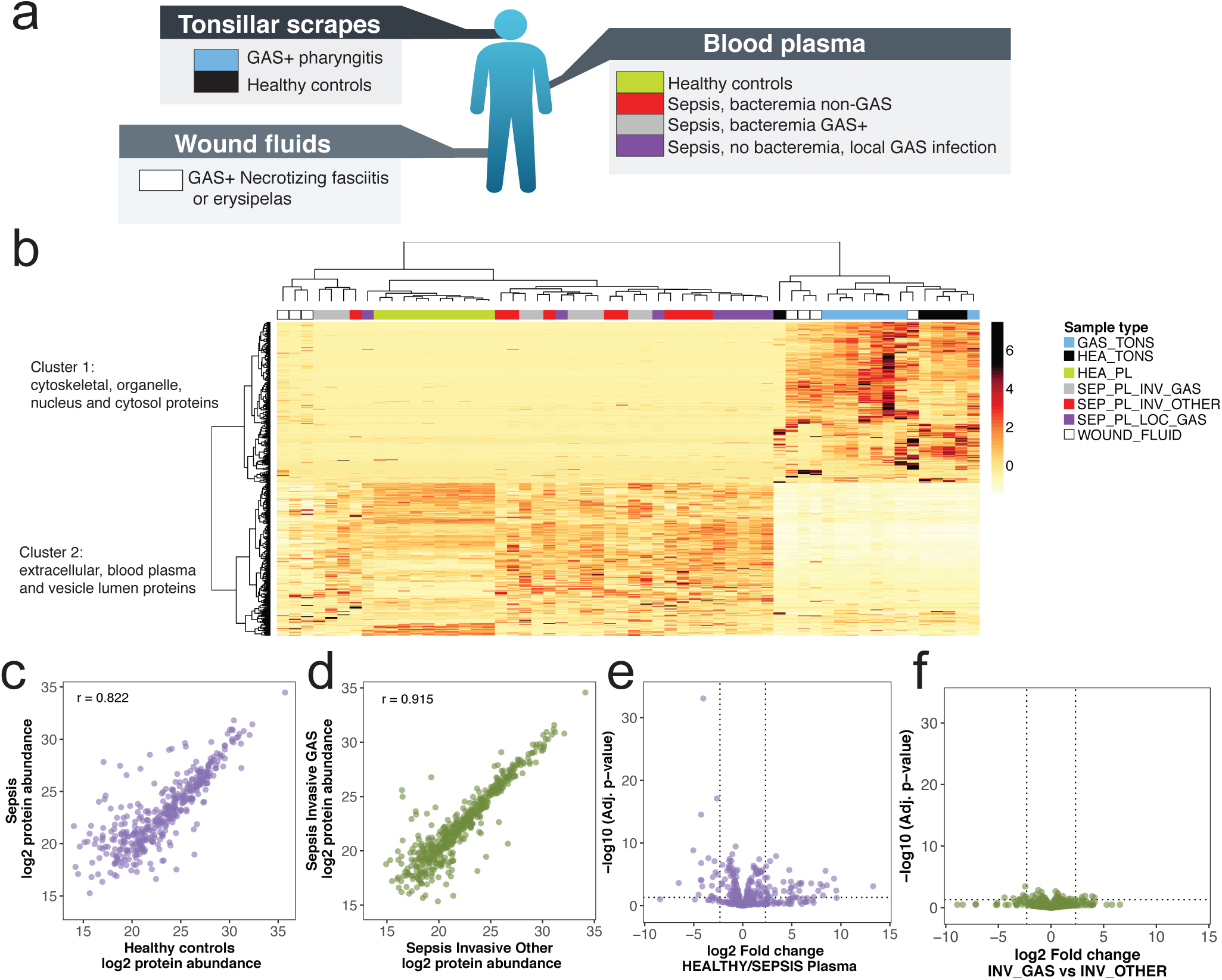
Proteomic analysis of human samples. A) Protein samples collected from patients diagnosed with local or systemic *S. pyogenes* infections or from healthy controls, were homogenized and digested with trypsin. Subsequent peptide samples were analyzed with shotgun and Data-independent acquisition (DIA) mass spectrometry-based proteomics workflows. The shotgun data was used for building a reference spectral library for all identified peptides and was used for targeted extraction of the DIA data. B) DIA data matrices were normalized by sample total intensities. The heat map shows the relative protein and protein group abundance quantified across samples. Unsupervised clustering using Pearson correlation delineates two major protein clusters and two major patient sample groups. Protein clusters are annotated by statistical overrepresentation test of Gene Ontology cellular component database. The patient samples are colours above based on patient group as same as in a) C-D) Correlation plots of protein median abundances from selected blood plasma sample groups and with Pearson correlation coefficient (r) indicated. E-F) Differential protein abundance volcano plots of selected blood plasma sample groups. Dotted lines indicate significance and fold change thresholds (vertically respectively horizontally). Multiple testing correction was performed by the Benjamini-Hochberg method.

### Differential levels of plasma proteins - a hall mark of infection

The incidence of pharyngotonsillitis caused by *S. pyogenes* is a 1000 fold higher compared to invasive *S. pyogenes* infections (Carapetis et al., 2005). The reason for this difference in incidence is not clear, but it has been shown that, for example, the IgGFc-binding surface proteins expressed by *S. pyogenes* protect the bacteria against phagocytic killing only in IgG poor microenvironments such as the skin and the oral cavity (Nordenfelt et al., 2012). Comparing the proteomes obtained from tonsil swabs from healthy controls and patients with *S. pyogenes* positive tonsillitis, revealed a significant increase of many proteins in the tonsillitis patients (Figure 4A). On average, the swabbing of tonsil surfaces in patients with tonsillitis has a 6-fold increase in protein mass compared to swabs from healthy controls (see Figure 4B and Table S4). The pairwise protein abundance t-tests with corrections for multiple testing, shows that the increase in proteome mass observed during tonsillitis is predominately attributed to infiltration of plasma proteins as indicated by the blue colored protein dots in the volcano plot in Figure 4A. The majority of the proteins with significantly increased protein levels are extracellular and/or exosomal proteins. In addition, a minor proportion of the increased proteins are associated with cellular structures such actin filament and glycolysis proteins (Table S4), possibly indicating tissue destruction or increased levels of cell infiltration. Next we analyzed the significantly differentially abundant tonsil proteins (p<0.05) across all sample types. Notably, the overlap between the significantly induced proteins in sepsis and tonsillitis is low (21 of 171/107 proteins respectively), demonstrating that the different disease conditions evoke different host responses. Clustering the significantly altered proteins in tonsillitis samples across all sample types provides a good separation of these samples and those from healthy controls into two distinct groups (Figure 4C). However, these proteins have no classification power in sepsis (Figure 4C, left sample cluster).

**Figure 4.**
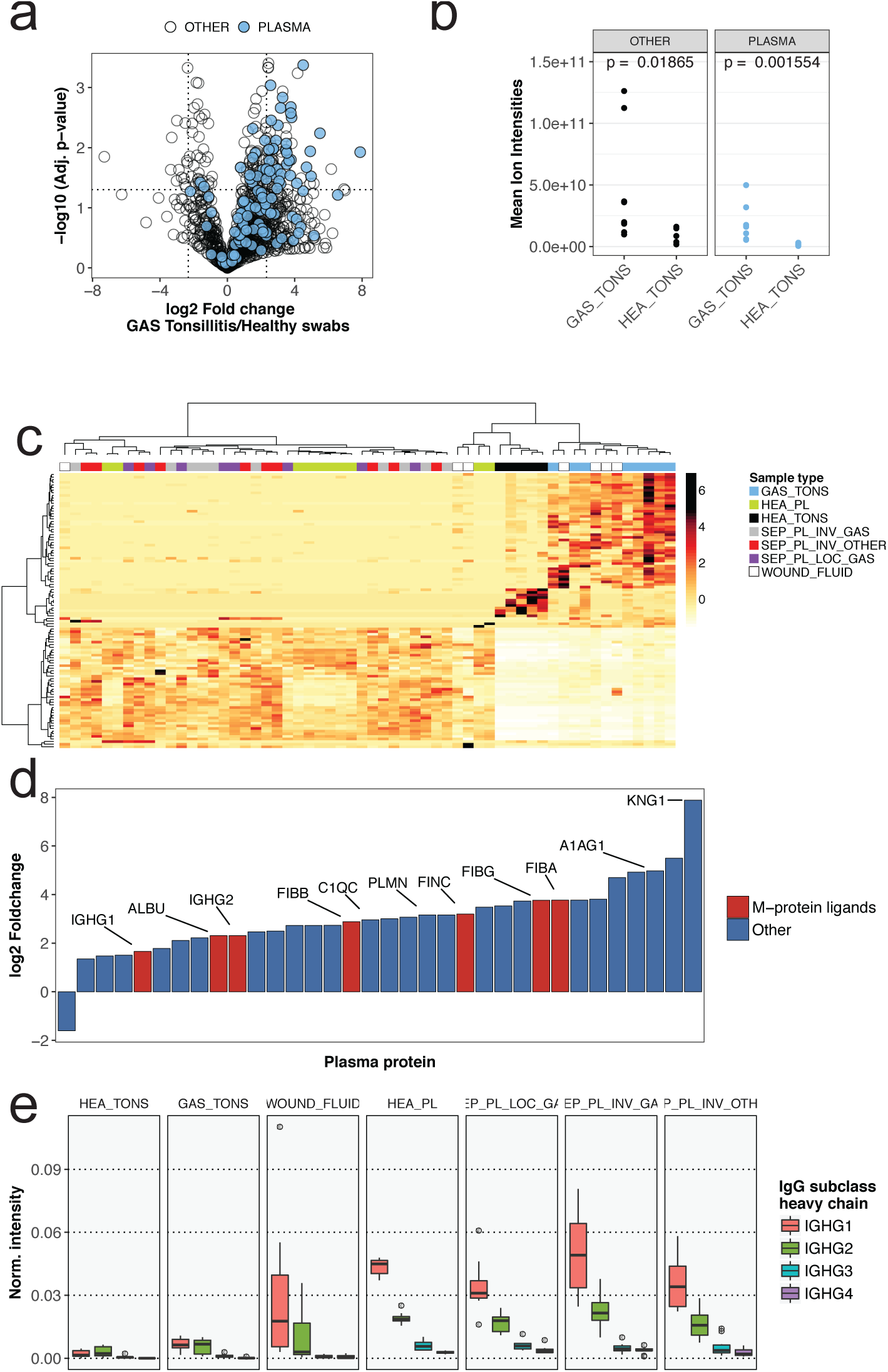
Differential abundance of IgG and other blood plasma proteins across patient and control samples. A) Differential protein abundance volcano plots of samples from tonsillar surfaces of *S. pyogenes* pharyngitis patients and healthy controls. Blue filled circles are blood plasma proteins. Dotted lines indicate significance and fold change thresholds (vertically respectively horizontally). Multiple testing corrections were performed by the Benjamini-Hochberg method. B) Comparison of mean total protein abundance (sum of all protein ion intensities) of tonsil swabs from patients with *S. pyogenes* pharyngitis (GAS_TONS) and healthy controls (HEA_TONS) showing plasma proteins (same as in a)) and other proteins. The P-values were calculated with non-paired Wilcoxon rank sum test. C) Hierarchical clustering of all patient samples with the subset of proteins that show significantly different abundance on healthy or *S. pyogenes* pharyngitis surfaces (Top left and right quadrants in Fig. 2a). D) Relative fold-change of plasma proteins with the significantly largest abundance difference between healthy and *S. pyogenes* pharyngitis surfaces (Top left and right quadrants in Fig. 2a). Plasma proteins with known binding to *S. pyogenes* M protein are filled with red, and others with blue. Lines and Uniprot Entry name highlight selected proteins with the common suffix “_HUMAN” removed. E) Normalized intensities (fraction of total signal) of IgG heavy chain subclasses across sample groups. Boxes represent 25^th^ to 75^th^ *%* quantiles, middle represents median, and lines smallest and largest observations.

Further analysis of the significantly increased plasma proteins during tonsillitis indicates that the plasma leakage is not uniform (Figure 4D). Based on the extracted ion intensities for the plasma proteins, we estimate an elevation of the plasma concentration at the site of infection from 16% to 28%. When including the increase in protein mass, the actual increase of plasma proteins during tonsillitis is in fact 10-fold (Figures 4A and B). Surprisingly, not all plasma proteins differed significantly between the groups (see Figure 4A), suggesting different propensity of individual plasma proteins to leak into the site of infection. The reason for this differential plasma leakage is not clear, but it is independent of plasma protein mass as there is no correlation between fold change and protein mass (data not shown). It is well established that *S. pyogenes* can bind numerous plasma proteins to its surface. The important *S. pyogenes* virulence factor M1 protein can for example bind fibrinogen (Kantor, 1965; Sanford et al., 1982), albumin (Åkesson et al., 1994; Myhre and Kronvall, 1980) and fibronectin (Schmidt et al., 1993) to inhibit phagocytosis (Carlsson et al., 2005) and facilitate uptake of fatty acids (J. A. Malmström et al., 2011). In our study, all known M1 protein interacting plasma proteins are among the top 30 proteins with the highest fold change during tonsillitis, as indicated by red bars in Figure 4D. The increased levels of these proteins at the site of infection may enable *S. pyogenes* to exploit these proteins to promote immune evasion. Alternatively, these proteins may be retained better at the site of infection possibly via their interaction with the streptococcal surface.

Since the efficiency of IgGFc-binding to M1 protein is dependent of IgG concentration, we investigated the IgG concentration in the different sample types. In the plasma samples the concentration of IgG is high, and sepsis does not introduce any major changes in IgG concentration or subclass distribution. The concentration of IgG is also on average high in wound fluid samples, although there is a considerable degree of variability in these samples. In contrast the tonsil swabs contain much lower amounts of IgG with higher levels of IgG during tonsillitis compared to the healthy controls (Figure 4D-E). Throughout all the analyzed samples we note that the subclass composition is not altered by *S. pyogenes.*

### IdeS activity in patients with *S. pyogenes* infection

In the following experiments we analyzed the samples outlined in Figure 3a (Table S1) using LC-SRM MS to quantify the absolute level of IgG cleavage in these samples (see Figure S5 and Table S5). Absolute quantification of IgG confirms that the IgG concentration is drastically different between site samples with high concentration of IgG in plasma samples and in some of the wound fluid samples (Figures 5A and B). These samples exhibit a concentration of around 1 prnol/μ! of IgG1, with an expected subclass distribution. The tonsil swabs have around 10 to 100-fold lower concentration of IgG compared to the plasma samples. The swabs from the tonsillitis patients have higher concentration of IgG compared to the swabs from the healthy volunteers, mostly pronounced for IgG1, due to increased plasma leakage as shown in Figure 4 This increase in plasma leakage was supported by the elevated levels of several other high abundant plasma proteins such as albumin and fibrinogen in the swabs from the tonsillitis patients (Figures 4B and 5A).

**Figure 5.**
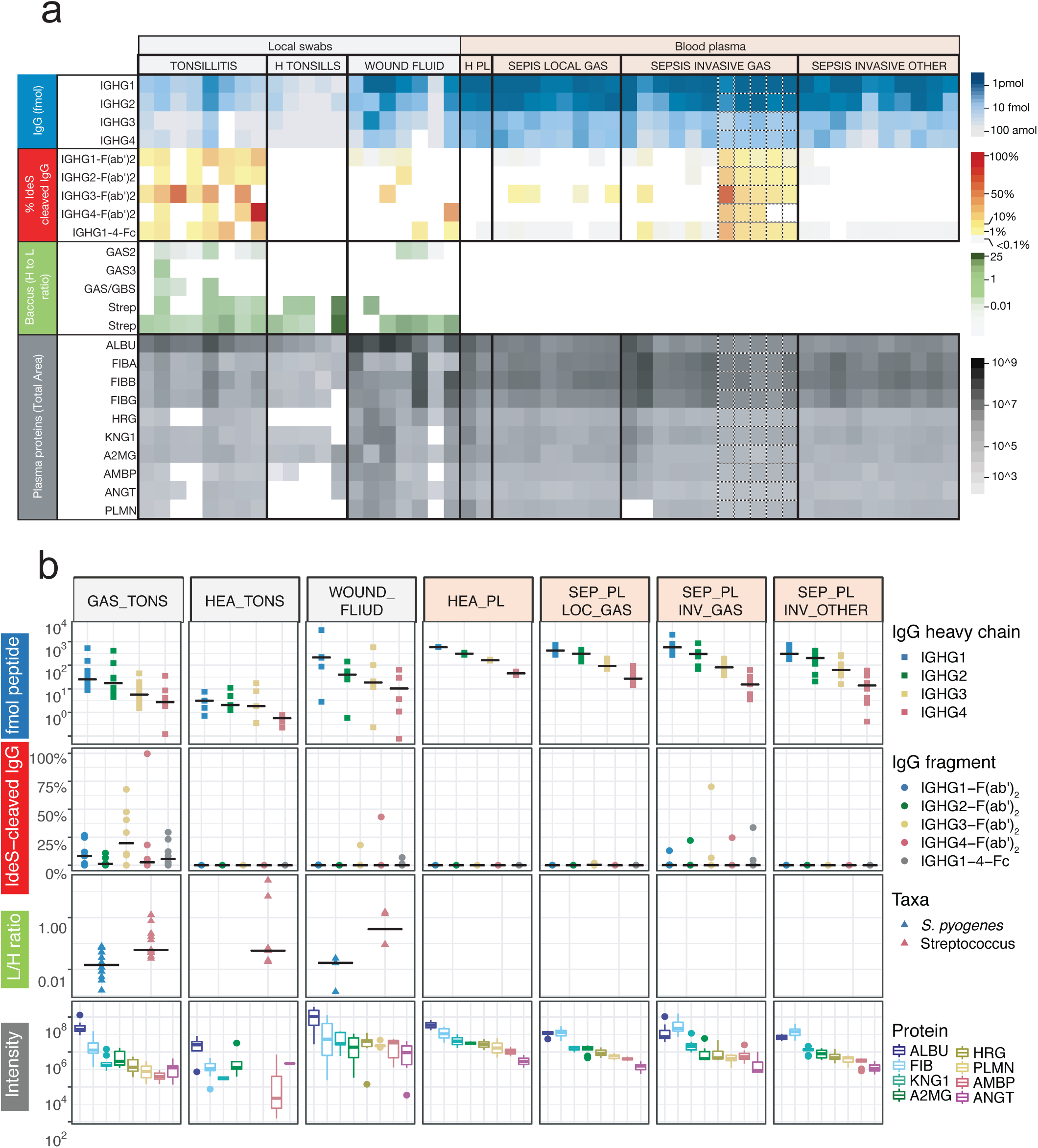
Analysis of IdeS-cleaved IgG and other targeted proteins in patient and control samples. The digested patient samples from Figure 3A were analyzed using the SRM-MS method show in Figure 1A. A) Each column represents a human sample collected either from a local infection, plasma from blood samples drawn from patients diagnosed with bacterial sepsis, or healthy control samples. Columns with dotted line borders are time series samples from a single patient (see Table S1). Blue panel shows the absolute concentration of total IgG. Red panel shows the proportion of IdeS-cleaved IgG. Green panel shows the levels of peptides specific for the *S. pyogenes* species or for the Streptococcus genus. Grey panel show the intensities of peptides derived from human plasma proteins. Abbreviations indicate the Uniprot Entry name with the common suffix “_HUMAN” removed. B) Shows the same data as in A), but by sample group and only the first time-point. Black lines indicate the group median value of y-axis values.

The tonsil swabs from tonsillitis patients have elevated levels of IgG, but still there is a considerable degree of IgG cleavage in these samples. All seven samples reveal signs of IdeS specific IgG cleavage, and in some cases specific subclasses are cleaved to 100%. Although IgG3 has a tendency to be more sensitive to IdeS cleavage, the differences between the subclasses are not significant. Surprisingly, we also observe detectable levels of cleaved IgG in the IgG rich samples such as some of the wound fluids and plasma sample from patients infected with *S. pyogenes,* but not in plasma from patients with sepsis caused by other bacterial pathogens. There are also trace levels of cleaved IgG in plasma from patients with sepsis with a confirmed local *S. pyogenes* infection but void of detectable levels of the bacterium in plasma determined using conventional culturing techniques. Notably peptides hypothetically derived from other defined IgG1 proteases were detected in several samples. However, the peptide intensities generally correlated to the total IgG1 levels (r = ≥ 0.55, p > 0.001). This was in contrast to IdeS-derived IgG1peptides which did not correlate with the total IgG1 heavy chain levels (r = 0.07, p = 0.73) (Figures S6A and S6B). In addition, the only IgG1 heavy chain peptide with large significant mean difference between *S. pyogenes* positive samples groups as compared to negative control groups, was the IdeS-derived peptide (Figure S6C). Combined these results demonstrate that IdeS cleavage of IgG either occurs locally or directly in the blood plasma and thereby noticeably alter the integrity of the complete IgG repertoire, and that the signals from IdeS cleavage IgG products are unique for samples from individuals infected by *S. pyogenes.*

To further explore the reason for the observed differential cleavage of IgG, we used a previously established *S. pyogenes* specific mass spectrometry based cell count method (Sjöholm et al., 2017). With this method we observe detectable levels of *S pyogenes* directly in three of the seven tonsillitis patients, with a pronounced signal in two patients. Similarly, the swabs from two of the three patients with necrotizing fasciitis revealed measurable levels of *S. pyogenes.* As expected, there were no detectable *S. pyogenes* specific peptides in the tonsil swab from healthy controls (see green panels in Figures 5A and 5B). In the majority of these samples we detect peptides specific for the *Streptococcus* genus reflecting the composition of the oral microbiota. Correlation analysis between the number of *S. pyogenes* cells and the degree of IgG cleavage reveal no correlation (data not shown). Surprisingly, we were unable to detect IdeS or SpeB in any of the samples and consequently it is not possible to correlate the degree of IgG cleavage to the level of IdeS or to determine if some of the *S. pyogenes* strains secrets higher concentration of IdeS. Furthermore, we are unable to rule out that the differences in cleavage may be attributed to neutralizing antibodies or other unknown factors influencing IdeS cleavage efficiency. Based on these results we conclude that lower absolute concentration of IgG and detectable levels of *S. pyogenes* cells are the predominant factors for proportionally high levels of IgG cleavage.

In conclusion, the novel SRM-MS assay identifies active IdeS-mediated degradation of IgG in all samples from patients with a confirmed *S. pyogenes* infection. At the local site of infection, we can quantify low levels of peptides unique for *S. pyogenes* although IdeS is present in amounts below the detection level. These results demonstrate that the *in vivo* activity of a low abundant and undetectable protease is capable of comprising part of the IgG pool with a higher cleavage efficiency in microenvironments associated with low IgG concentrations.

## Discussion

The aim of the present study was to investigate how the human proteome is influenced by *S. pyogenes* infections, locally and during invasive disease. To address this question a novel mass spectrometry approach was developed and applied to a unique set of samples from patients and healthy individuals. A combined quantitative and targeted mass spectrometry strategy was used, and the patient material included both swabs from local infections and sepsis blood plasma and wound fluids from patients with invasive disease.

Sepsis results in a dramatic change in the blood plasma proteome composition as compared to healthy conditions. Some of the proteins with significant changes are known markers for inflammation (acute phase proteins) and are expected to increase or decrease due to the disease. However, when comparing sepsis caused by *S. pyogenes* with other bacterial pathogens the difference was minor. Sepsis is a serious and lifethreatening condition but only constitutes a minor part of all *S. pyogenes* infections. *S. pyogenes* most common habitat is the human pharynx, where it either causes infection or resides asymptomatically. In asymptomatic carriers, the bacteria are exposed to low levels of blood plasma proteins originating from saliva and mucosal secretions, whereas during tonsillitis/pharyngitis, the inflammation will cause exudation of plasma proteins to the epithelial surface. Here we detect a 10-fold increase of many plasma proteins in the swabs from patients with *S. pyogenes* tonsillitis, including plasma proteins known to interact with *S. pyogenes* surface proteins. Since the increase of individual plasma proteins is not dependent on molecular mass, the pattern is not explained simply by plasma leakage, but rather by the binding and retainment of proteins at the surface of infecting *S. pyogenes* bacteria.

Based on the novel SRM-based IgG quantification assay, we could confirm for the first time that IdeS hydrolyses IgG during a human *S. pyogenes* infection, and that the cleavage efficiency is highly variable depending on the site of infection. On average, the highest levels of IdeS-cleaved IgG are found in tonsillitis samples, much higher than in plasma from sepsis patients. This is expected considering the lower concentrations of IgG at the site of a local infection compared to vascular IgG levels. The consequences of IdeS activity on IgG immune responses have been extensively described and discussed previously (Pawel-Rammingen, 2012; Pawel-Rammingen et al., 2002b; 2002a), and in this work we find that the IgG3 subclass is most susceptible to IdeS cleavage *in vitro,* as well as in the IdeS clinical trial on healthy volunteers and during *S. pyogenes* infection *in vivo.* IgG3 has been proposed to be the first IgG subclass to emerge after IgM isotype switching and involved in the early humoral response against pathogens (Collins and Jackson, 2013). Moreover, IgG3 has the highest affinity for complement factor C1q (Schroeder and Cavacini, 2010). Hypothetically, IdeS has evolved primarily to target IgG3 functions, but the results could also be due to IgG3’s inherent lower half-life and increased susceptibility to hinge-region proteolysis (Carter, 2006).

When recombinant IdeS is intravenously administered to healthy humans, the total IgG pool is rapidly cleaved (Winstedt et al., 2015) (Fig. 2). However, in this case the concentration of IdeS (~6 μg/ml) is orders of magnitude higher than what can be expected during an infection with *S. pyogenes.* To exemplify; the mass of a streptococcal cell (dry weight) is 0.2 pg of which 55% is protein. With a high bacterial load of 10^3^ bacteria per ml blood, this corresponds to 110 pg of *S. pyogenes* total protein per ml blood. Although the enzyme and substrate dynamics are different during an infection with IdeS production over time and perhaps reaching a steady-state concentration of the protease, the concentration will be substantially lower than what is required for the cleavage of the entire IgG pool. IdeS is a uniquely specific protease, apart from IgG no other substrate has been identified, and based on current knowledge, the cleavage sites in IgG heavy chains are also unique for IdeS (Brezski and R. E. Jordan, 2010). The present work shows that IdeS activity above background levels (>0.1% cleavage) strongly correlates with *S. pyogenes* infection. It is most likely that IdeS is present during *S. pyogenes* infection since the *ideS* gene is present in most serotypes of *S. pyogenes* (Pawel-Rammingen, 2012) and serum samples from individuals following *S. pyogenes* infection contain anti-IdeS antibodies (Åkesson et al., 2006). Furthermore, screening of 208 serum samples from healthy humans showed that >95% have detectable levels of anti-IdeS antibodies (Winstedt et al., 2015), showing that most humans have been infected with IdeS-producing *S. pyogenes* stains during their lifetime. There are several other bacterial and eukaryotic proteases that hydrolyze IgG in the hinge region, but not at the same residual sites as IdeS (Brezski and R. E. Jordan, 2010). However, it cannot be excluded that so far uncharacterized IdeS-like enzymes might be produced by other members of the microbiota at epithelial surfaces colonized by *S. pyogenes,* or that IdeS-like Fc and F(ab)_2_ fragments could be generated by non-specific cleavage by broad-spectrum human proteases released as part of the inflammatory response to *S. pyogenes* infection.

The SRM-method estimates the percentage of IdeS-cleaved IgG based on fragment-ion relative intensities of the light and heavy isotope-labelled versions of absolutely quantified peptides. Heavy labelled peptides represent the gold-standard for peptide quantification, although this quantification has its limitations and drawbacks (Picotti and Aebersold, 2012). The heavy peptides do not account for differences in the proteolytic cleavage efficacy or peptide loss in the sample preparation step. Hence the relationship between inherent peptide and the original protein levels may not be equimolar (Abbatiello et al., 2015). This effect is likely amplified in our study as our method requires two peptide pairs to calculate the amounts of IdeS-cleaved IgG. Normally SRM-based quantification involves selection of optimal peptides to represent their parent protein. When measuring IdeS activity on IgG as performed here, the target peptides were predetermined by the specificity of IdeS cleavage of the lower hinge region of IgG, which gives no option to discard suboptimal peptides. We tried to circumvent these limitations by making calibrations curves with known levels of intact and cleaved IgG. However, this indirect mode of measuring IdeS activity is likely associated with errors in the quantification. Previous method of measuring IdeS activity is performed by for example densitometric measurements of stained bands on SDS-PAGE gels or ELISA assays. Still, the SRM method developed here is clearly associated with improved quantitative accuracy and dynamic range, and, which is important in relation to this study, the method offers the possibility of measuring several related aspects such as levels of IdeS, absolute amount of IgG and other plasma proteins, and additional cleavage patterns known to compromise the structure of IgG.

This work provides a novel experimental approach to investigate the effect of a significant bacterial pathogen on the human proteome in different clinical conditions; asymptomatic colonization, localised superficial infection and invasive disease. In addition, the analysis of the IgG degrading activity of IdeS in healthy individuals and patients infected with *S. pyogenes*, shows that the specific activity of a bacterial protease can be used to diagnose and follow/evaluate the progression of an ongoing infection and also makes IdeS a promising vaccine candidate. This provides new scientific opportunities with considerable clinical and theoretical potential.

## Material Methods

### Synthetic peptides and purified proteins

Non-purified synthetic peptides were purchased from JPT Peptide Technologies or ProteoGenix. Heavily labelled peptides were purchased from ThermoFisher Scientific (AQUA-grade QuantPro). For further details on synthetic and labelled peptides see Table S2. Recombinant and quantified IdeS was from Hansa Medical AB (Winstedt et al., 2015). Human monoclonal IgG subclasses were purchased from Sigma-Aldrich.

### Human samples

Serum samples from healthy individuals treated with IdeS were from the Phase I trial NCT01802697 (Winstedt et al., 2015). Briefly, subjects were intravenously injected with IdeS (at the following doses: 0.01, 0.04, 0.12 or 0.24 mg IdeS/ kg bodyweight), followed by a time-course collection of serum samples. Plasma samples from septic patients were from Linder et al., 2009. Plasma samples from healthy individuals were from Stenemo et al., 2016 or purchased from Innovative Research (pooled normal human plasma). Swabs and scrapes from local infections from patients with diagnosed *S. pyogenes* infection were collected at Skåne University Hospital or at Laurentiikliniken (a primary health clinic), both in Lund, Sweden. Tonsillar swabs from healthy individuals were obtained at Lund University, Lund Sweden. The medical ethics committee of Lund University approved this study (protocol numbers 2005/790 and 2015/14) and informed consent was obtained from all subjects. The samples were transported at −20°C and stored at - 80°C until further processing. A compilation of patients, control and IdeS clinical phase I study samples is in Table S1.

### IdeS endopeptidase assays

66 pmol of respective purified monoclonal IgG subclass was mixed with 1 pmol IdeS and incubated at 37°C. Aliquots were removed at time-points 1, 10 and 120 minutes and enzymatic activity was stopped by denaturing with 8 M urea. For time-points 0 min, IdeS and IgG were mixed in 8 M urea directly. The samples were analysed with nonreducing SDS-PAGE Criterion TGX Gels (Bio-Rad) or SRM (after tryptic digestion, see below). Intact and IdeS-cleaved IgG was calculated by summarizing gel-band intensities. IdeS activity on SDS-PAGE was quantified densitometrically using software GelAnalyzer 2010a. Correlations between SRM and SDS-PAGE were done with linear model methods using the software GraphPad Prism 6.

### Sample preparation

Swab and scrape samples were homogenized using a Fastprep-96 beadbeater (MPBio) with Lysing Matrix B tubes (MPBio). All protein samples were denatured with 8 M Urea in 100 mM ammonium bicarbonate (ABC) and then reduced with Tris (2-carboxyethyl) phosphine (TCEP) (Sigma-Aldrich) for 1 h at 37°C, and alkylated with 25 mM iodoacetamide for 45 min before diluting the sample with 100 mM ABC to a final urea concentration below 1.5 M. Proteins were digested by incubation with trypsin (1/100, w/w) for 10 h at 37°C. The peptides were desalted and cleaned-up with reversed-phase spin columns (Vydac UltraMicroSpin Silica C18 300Å Columns, Harvard Apparatus) according to the manufacturer’s instructions.

### Data dependent acquisition (DDA) and Data independent acquisition (DIA)

These peptide analyses were performed on a Q Exactive Plus mass spectrometer (Thermo Scientific) connected to an EASY-nLC 1000 ultra-high-performance liquid chromatography system (Thermo Scientific). Peptides were separated on an EASY-Spray ES802 columns (Thermo Scientific) using a linear gradient from 3 to 35% acetonitrile in aqueous 0.1% formic acid during 2 h. DDA and DIA instrument settings were identical to as described by (E. Malmström et al., 2016). From the DDA data a spectral library was built using the workflow described by (Schubert et al., 2015) using the Trans-proteomic pipeline (TPP v4.7 POLAR VORTEX rev 0) and Human UniProtKB/Swiss-Prot release 2015_10 canonical sequences, synthetic peptide sequences (Table S2) or *S. pyogenes* NCBI reference sequence NC_002737.1. The human generated spectral library was used to extract the DIA data with the DIANA algorithm v2.0.0 (Teleman et al., 2015) using a 1% peptide false discovery rate. For protein quantification, the top3 integrated peptide ion intensities extracted from the MS2 spectra was summed up by protein or protein group (Ahrné et al., 2013).

### Selected reaction monitoring (SRM)

The SRM measurements were performed on either TSQ Vantage or TSQ Quantiva triple quadrupole mass spectrometers (Thermo Scientific), connected to EASY-nLC II high-performance liquid chromatography systems (Thermo Scientific). Peptides were separated on columns PicoChip PCH7515-105H354-FS25 (New Objective) or EasySpray ES800 (Thermo Scientific), using a linear gradient from 5 to 35% acetonitrile in aqueous 0.1% formic acid during 34 min. In general, instrument dwell time settings were fixed at 10 ms or 40 ms for high respectively low signal intensity peptides, and at scheduled acquisition times defined by iRT normalization (Escher et al., 2012). SRM assays were obtained from the following publications (Karlsson et al., 2012; Nordenfelt et al., 2012; Sjöholm et al., 2017; 2014), or developed as described by (Lange et al., 2008). The raw data was processed and analyzed with SRM analysis software Skyline (MacLean et al., 2010), with manual validation and inspection of the results. This included signal intensity, relative fragment ion intensities ratios based on DDA spectral libraries and retention times.

### Data analysis and statistics

The PANTHER database (Mi et al., 2013) release 20160715 was used for Overrepresentation Test with GO cellular component complete database using Bonferroni correction for multiple testing. Applied data analysis was done using custom R scripts, mainly using tidyverse, reshape2, aHeatmap, and ggpubr packages.

## Acknowledgments

We thank personnel at Laurentiikliniken, Lund, Sweden, and Skåne University Hospital, Lund, Sweden, for help with patient sample collection; S. Hauri and J. Teleman for providing MS data processing shell scripts; and A. Nägeli and S. Hauri for help with sample preparation and MS analysis.

This work was supported by the Foundations of Knut and Alice Wallenberg, Torsten Söderberg and Alfred Österlund, Olle Engkvist Byggmästare, Crafoord (grant 20100892) as well as the European research council starting grant (ERC-2012-StG-309831), the Swedish Research Council (projects 7480 and 621-2012-3559), the Wallenberg Academy Fellow program KAW (2012.0178 and 2017.0271) the Swedish Government Funds for Clinical Research (ALF), and the Medical Faculty of Lund University.

## Author Contributions

Conceptualization, C.A.Q.K, A.L and, J.A.M; Methodology, C.A.Q.K, A.L and, J.A.M; Investigation C.A.Q.K, S.J., L.W., C.K., L.B., A.L. and J.A.M.; Writing – Original Draft, C.A.Q.K and J.A.M; Writing – Review & Editing, C.A.Q.K, S.J., L.W., C.K., L.B., A.L. and J.A.M.; Funding Acquisition, L.B., A.L. and J.A.M; Supervision, C.K., L.B., A.L. and J.A.M.

## Supplemental Information

Figures S1-S6 and Tables S1-S5.

The MS raw data and applied analysis results are deposited at PeptideAtlas (ftp://PASS01096:DG4298dm@ftp.peptideatlas.org/) and Panorama https://panoramaweb.org/labkey/xanZMv.url, user, password: rmrcb6mxY).

